# Cargo crowding drives sorting stringency in COPII vesicles

**DOI:** 10.1101/2020.03.04.976183

**Authors:** Natalia Gomez-Navarro, Alejandro Melero, Jerome Boulanger, Wanda Kukulski, Elizabeth A. Miller

**Affiliations:** MRC Laboratory of Molecular Biology

## Abstract

Accurate maintenance of organelle identity in the secretory pathway relies on retention and retrieval of resident proteins. In the endoplasmic reticulum (ER), secretory proteins are packaged into COPII vesicles that largely exclude ER residents and misfolded proteins by mechanisms that remain unresolved. Here we combined biochemistry and genetics with correlative light and electron microscopy (CLEM) to explore how selectivity is achieved. Our data suggest that vesicle occupancy dictates ER retention: in the absence of abundant cargo, non-specific bulk flow increases. We demonstrate that ER leakage is influenced by vesicle size and cargo occupancy: overexpressing an inert cargo protein, or reducing vesicle size restores sorting stringency. We propose that cargo recruitment into vesicles creates lumenal steric pressure that drives selectivity. Sorting stringency is thus an emergent property of the biophysical process of cargo enrichment into a constrained spherical membrane-bound carrier.

**Summary:** Combining correlative light and electron microscopy with yeast genetics and biochemistry, Gomez-Navarro, Melero and colleagues show that cargo recruitment into a constrained COPII vesicle restricts bulk flow, thereby contributing to sorting stringency and ER quality control.

## Introduction

Protein trafficking within the eukaryotic secretory pathway occurs via cargo-bearing vesicles that shuttle proteins and lipids from one compartment to another. Cytosolic coat proteins drive vesicle formation by deforming the membrane of the donor organelle into small carriers and selecting cargo proteins for incorporation into the carrier vesicles (for reviews see (Bonifacino and Lippincott-Schwartz, 2003; Dancourt and Barlowe, 2010; Geva and Schuldiner, 2014)). The first step taken by nascent secretory proteins is packaging into COPII-coated vesicles that bud from the endoplasmic reticulum for delivery to the Golgi (Barlowe et al., 1994; Gürkan et al., 2006; Lee et al., 2004). The COPII coat assembles on the ER membrane in two layers. The inner cargo- and lipid-bound layer comprises the small GTPase, Sar1, and the cargo adaptor complex, Sec23/Sec24. This inner coat in turn recruits an outer coat of heterotetrameric Sec13/Sec31, which forms rod-like structures that can self-assemble into a polyhedral cage that is thought to contribute to vesicle architecture (Fath et al., 2007; Noble et al., 2013; Zanetti et al., 2013). In addition to the five core COPII coat proteins, regulatory components control vesicle formation at discrete ER exit sites (ERES). Sec16 is one example of an accessory protein that is thought to define sites for COPII recruitment and assist in coat assembly (Supek et al., 2002; Kung et al., 2012).

ER exit can be highly selective: in some cell types and in *in vitro* reconstitution experiments, properly folded secretory proteins are enriched in COPII vesicles, and ER resident proteins are largely excluded (Barlowe et al., 1994). Indeed, despite the very high concentrations of ER resident proteins (Macer and Koch, 1988), secretion of ER chaperones and folding intermediates is minimal. Cargo enrichment into COPII vesicles is mediated by direct interaction between ER export signals and Sec24, which contains multiple independent cargo-binding sites (Miller et al., 2003; Mossessova et al., 2003; Mancias and Goldberg, 2007; 2008). Protein sorting is also facilitated by cargo receptors that bridge the interaction between cargo and coat proteins (Geva and Schuldiner, 2014). In addition to signal-mediated trafficking, proteins can also move within the secretory pathway by bulk flow, whereby proteins are not enriched in vesicles but are stochastically captured at their prevailing concentrations as part of the bulk fluid or membrane (Martínez-Menárguez et al., 1999; Wieland et al., 1987; Polishchuk et al., 2003; Thor et al., 2009).

An abundant class of cargo proteins, the GPI-anchored proteins (GPI-APs), are packaged into COPII vesicles via interaction with the p24 family of proteins (Castillon et al., 2011). Deletion of any of the four major yeast p24 proteins (Emp24, Erv25, Erp1 and Erp2) results in multiple phenotypes: defective ER retention of both resident and misfolded proteins; activation of the Unfolded Protein Response (UPR); and viability in the absence of Sec13 (known as a *bst,* for <u>b</u>ypass of <u>s</u>ec-<u>t</u>hirteen, phenotype) (Elrod-Erickson and Kaiser, 1996; Belden and Barlowe, 2001; D’Arcangelo et al., 2015). The molecular basis for these phenotypes remains poorly understood, including how the different cellular outcomes relate to each other. One model for the *bst* phenotype is that concentration of GPI-APs at ERES creates a local domain that is resistant to membrane deformation (Copic et al., 2012; D’Arcangelo et al., 2015). This rigid membrane requires the COPII coat to do extra work to enforce curvature, work that is contributed in part by Sec13. Thus, in p24 mutants, where GPI-AP enrichment is reduced, the absence of Sec13 is tolerated because less force is required to overcome the membrane bending energy at an ERES. Whether the ER leakage and UPR phenotypes are also related to membrane rigidity and coat scaffolding is unknown.

In addition to their function as GPI-AP receptors, p24 proteins have also been postulated to function as a quality control filter by modulating the timing of vesicle release (Kaiser, 2000), or by displacing non-specific cargo (Kaiser, 2000; Ma et al., 2017). Here we aimed to better understand the consequences of p24 deletion, focusing on two aspects: membrane bending by Sec31 alone, and selectivity of ER export. We examined vesicle morphology *in situ* in the absence of Sec13, and probed mechanisms by which the p24 proteins might act as a selectivity filter. We find that in the absence of Sec13, COPII-associated membranes become large and pleiomorphic. In contrast, ER leakage in the *emp24Δ* mutant derives from alterations in cargo occupancy rather than vesicle morphology or specific p24 function. We propose that inclusion of p24 proteins and other abundant cargoes in COPII vesicles creates steric effects that preclude inappropriate capture of ER residents.

## Results

To understand the basis for vesicle formation in the absence of Sec13, we first sought to visualize the ultrastructure of ERES with high spatial resolution. Sites of COPII vesicle formation can be localised in cells expressing fluorescently tagged COPII subunits (Watson et al., 2006; Okamoto et al., 2012). However, membrane morphology falls below the diffraction limit such that detailed structural information can only be obtained by electron microscopy (Orci et al., 1991; Zeuschner et al., 2006). To exploit the advantages of fluorescent protein localisation and the resolution of electron tomography, we used a correlative light and electron microscopy (CLEM) approach (Kukulski et al., 2011). We tagged Sec24 at its chromosomal locus by integrating superfolder GFP (sfGFP) at the C-terminus (Sec24-sfGFP) and subjected cells to high pressure freezing, freeze substitution and resin embedding. Thick sections (∼300nm) were collected on EM grids, labelled with fluorescent fiducial markers and imaged by fluorescence microscopy to identify GFP-positive ERES (Figure 1A, inset). Sections were subsequently imaged by electron tomography at both low (Figure 1A, left panel) and high magnification (Figure 1A, centre and right panels). Fiducial markers permit the precise spatial correlation of the Sec24-sfGFP signal within the electron tomograms, allowing visualization of the underlying membranes in this region (Figure 1A, right panels). We found a range of membrane morphologies at Sec24-positive ERES, including flat ER membranes, budding events with a nascent vesicle still continuous with the ER, ERES with multiple buds, and free vesicles released from ER membranes (Figure 1A, 1B and S1A).

**Figure 1.**
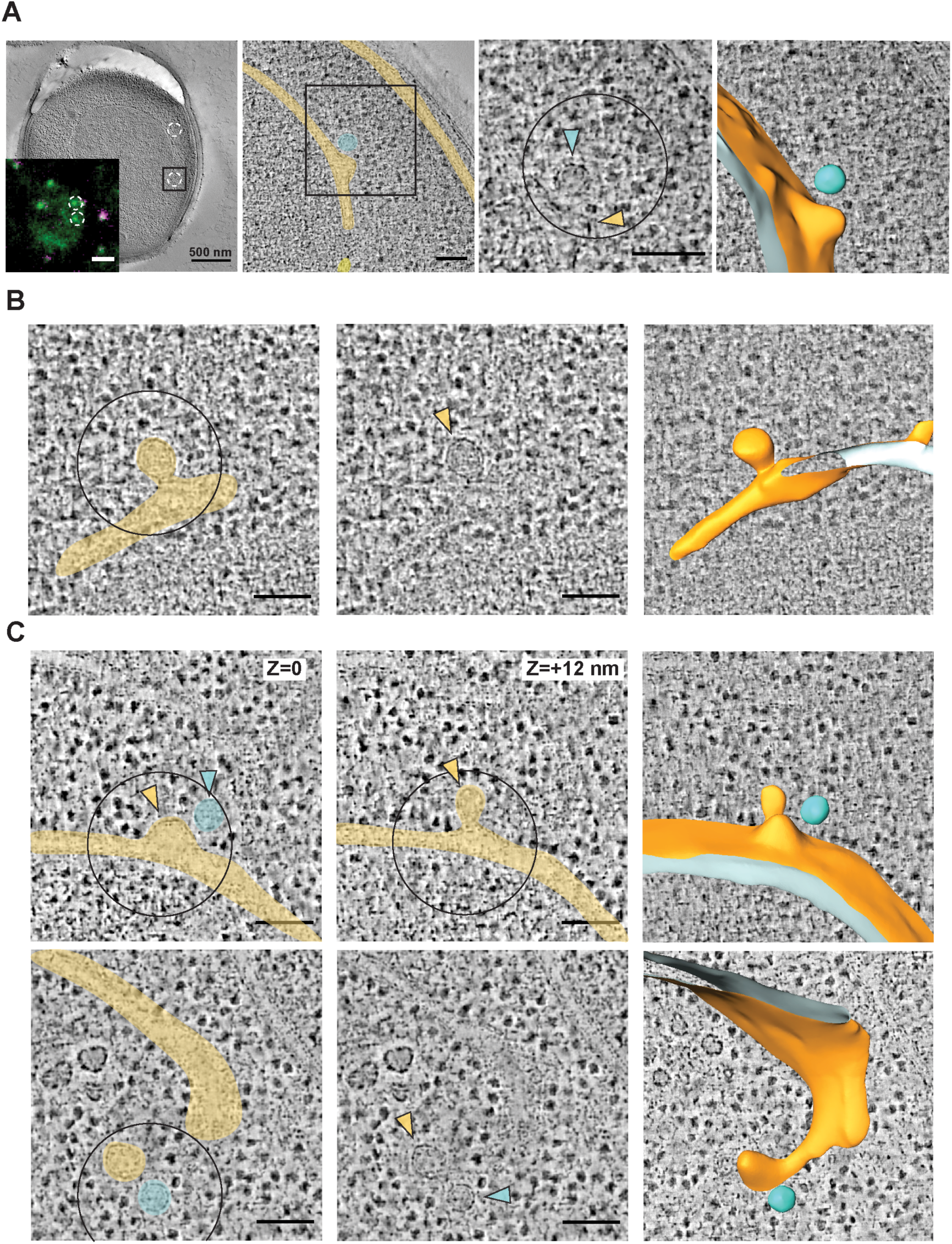
3D ultrastructure of COPII budding events and free vesicles by CLEM. (A) Resin-embedded yeast cell containing ERES correlated to Sec24-sfGFP signal. Left panel, inset: two Sec24-sfGFP foci (green) are highlighted. Fiducial markers used for correlation are visible in green and magenta. Left panel, a virtual slice of a low magnification tomogram used for correlation with fluorescent image of the same cell (inset). Middle-left panel is a virtual slice of a high magnification tomogram on one of the two Sec24-sfGFP marked ERES, highlighted by a 500 nm^2^ square. Nuclear envelope and ER cisterna are false-coloured in yellow and a free vesicle is highlighted in cyan. Middle-right panel is a zoom-in on the same virtual slice as previous panel. The centre of the 250 nm diameter circle marks the predicted position of the GFP signal centroid. Coloured arrowheads mark a nascent COPII bud (yellow) and a free-standing vesicle (cyan), represented in a segmentation model on the farthest right panel. (B) A virtual tomographic slice showing a correlated Sec24-sfGFP ERES. Left panel, ER is false-coloured in yellow. Central panel, arrowhead marks the emerging bud. Right panel is a segmentation model of the budding event. (C) Upper panels: two virtual tomographic slices of a single Sec16-sfGFP correlated spot: different z positions of the same x,y position are shown, revealing a multibudded ERES with 2 buds (yellow) and a free vesicle (cyan). Lower panels are a virtual tomographic slice showing a correlated Sec16-sfGFP ERES. Left panel, ER is false-coloured in yellow. Central panel, arrowheads mark an emerging bud (yellow) and a free-standing vesicle (cyan). Right panel is a segmentation model of the budding event. Scale bars A: EM, 500 nm, inset, 1 µm,; all other panels: 100 nm.

We also introduced sfGFP at the chromosomal locus of *SEC16* of wild-type cells, revealing similar ERES as Sec24-sfGFP (Figure S1C). By CLEM we observed similar membrane morphologies associated with both Sec16-sfGFP (Figure 1C) and Sec24-sfGFP (Figures 1A, 1B and S1A). Together, Sec24-sfGFP and Sec16-sfGFP strains yielded a combined dataset of 127 electron tomographic reconstructions of sites of COPII vesicle formation.

### ERES are pleiomorphic in the absence of Sec13

To visualize COPII budding events in the absence of Sec13 we attempted to chromosomally GFP-tag Sec24 in the *sec13Δ emp24Δ* strain, but were unsuccessful, suggesting that a tag on Sec24 is not well tolerated in this background (Figure S1B). Instead, Sec16-sfGFP in this mutant background showed viability similar to that of the *sec13Δ emp24Δ* parental strain (Figure S1B) and ERES similar to those of Sec24-sfGFP in wild type cells (Figure S1C).

Having validated that similar structures can be observed with Sec16-sfGFP as with Sec24-sfGFP in wild type cells, we next applied CLEM to *sec13Δ emp24Δ SEC16-sfGFP* cells. In the absence of Sec13, the N-terminal β-propeller region of Sec31 is thought to remain intact and capable of assembling into a polymeric cage (Fath et al., 2007; Stagg et al., 2006; Copic et al., 2012). We anticipated that the flexible hinge region of Sec31 exposed by loss of Sec13 should yield a less rigid structure with reduced membrane bending capacity, thereby generating enlarged COPII buds and vesicles. We acquired 31 tomograms at Sec16-sfGFP-correlated ERES in *sec13Δ emp24Δ* cells (Figures 2A and S2). The corresponding membrane ultrastructures showed broadly similar characteristics as those in wild-type cells, encompassing flat ER membranes, budded and multi-budded structures. However, the sites in *sec13Δ emp24Δ* cells appeared more pleomorphic, with numerous buds and vesicles (Figures 2A and S2).

**Figure 2.**
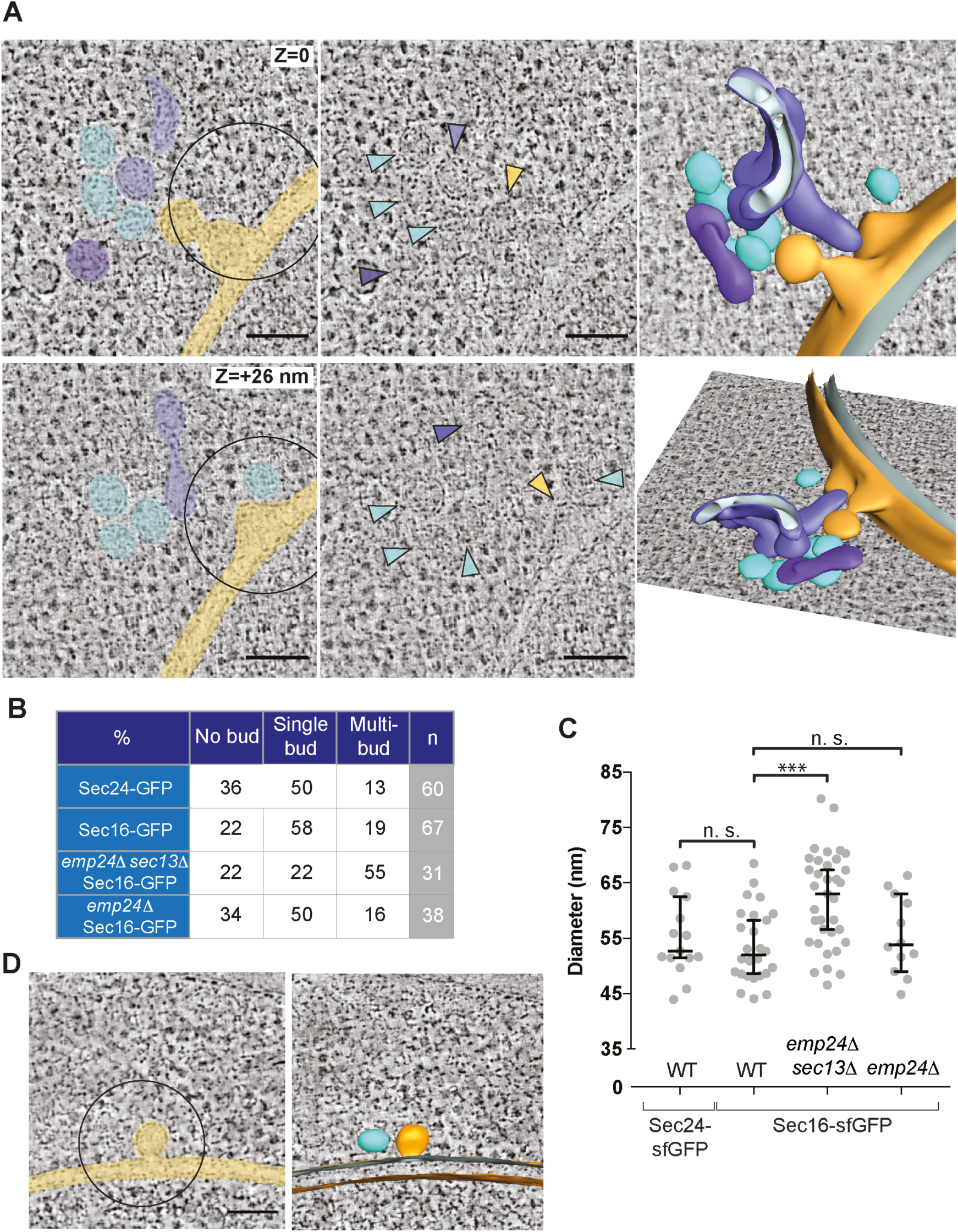
Deletion of *SEC13* results in pleomorphic membranes at ERES. (A) *SEC16-sfGFP-positive* ERES localised by CLEM in an *emp24*Δ *sec13*Δ cell. Upper and lower panels are different virtual slices from the same tomogram, representing different z-positions. Two buds form at the nuclear envelope (yellow) with 6 free vesicles (cyan) and 2 undefined tubular compartments (purple) in close proximity. In the central panels the same membrane structures are marked by coloured arrowheads. Right panels are two side-views of a segmentation model of the ERES. (B) Table of ERES ultrastructure categories (percentages from total number (n) of correlated spots per yeast strain). (C) Plot of maximum diameter (nm) of vesicles. Bars correspond to median value and 95% confidence interval. Statistical test was a t-test, *** p<0.0001. (D) A *SEC16-sfGFP-positive* ERES localised by CLEM in an *emp24*Δ *SEC16-sfGFP* cell. A bud emerges from the nuclear envelope (yellow) with a free vesicle by the side (cyan). 3D segmentation model on the right. Scale bars, 100 nm.

To quantify the different morphologies of ERES, we designated three broad categories of structures: flat ER membranes without an obvious bud (no bud), ER membranes with a single budded structure still attached (single bud); and structures with multiple curved membrane structures (multi bud) (Figure 2B). Comparison of the classes from different strains revealed no obvious difference between wild-type ERES marked with either Sec24-sfGFP or Sec16-sfGFP. Most ERES in both wild-type strains showed a single budding event, while multi-budded structures were less frequent. In contrast, the *sec13Δ emp24Δ* strain had fewer single bud profiles and many more multi-budded membranes (Figure 2B). As a second measure of membrane morphology, we used a quantitative segmentation analysis to measure the sizes of free vesicles (Machado et al., 2019). Maximum diameters of vesicles ranged from 45 nm to 65 nm, with a median of 52 nm (Figure 2C). Vesicles in *sec13Δ emp24Δ* cells showed a broader size distribution than in wild-type cells, with a median diameter 18% larger than wild-type (Figure 2C). We note that the smallest wild-type vesicle diameters of 45-50 nm were also found in *sec13*Δ *emp24*Δ cells, suggesting that Sec13 is not absolutely required to achieve high curvature (Figure 2C).

The large pleiomorphic structures of COPII-associated membranes in the *sec13Δ emp24Δ* strain suggest an obvious mechanism of lax ER retention, one of the shared phenotypes of the p24 mutants. When larger structures bud from the ER, bulk flow capture would increase, thereby stochastically packaging more ER residents and misfolded nascent proteins. This could overwhelm Golgi-ER retrieval mechanisms and result in leakage of ER residents to the cell surface, generally measured as secretion of the lumenal HSP70, Kar2 (Semenza et al., 1990; Schuldiner et al., 2005). We examined the morphology of Sec16-sfGFP-associated ERES in the *emp24Δ* single mutant to determine if large membrane structures might explain ER retention defects in single mutant cells. However, we observed largely normal membranes: in 38 tomograms of Sec16-sfGFP-positive regions (Figure 2D), most had the simple ERES morphology observed in wild-type cells comprising a single nascent bud (Figure 2B). Multi-budded structures were less common in the *emp24Δ* single mutant than in the *sec13Δ emp24Δ* double mutant, and were observed at a similar frequency as in wild-type cells. Furthermore, wild-type cells and the *emp24Δ* single mutant showed similar size distributions and median diameters of free vesicles (45 nm to 65 nm, median of 54 nm) (Figure 2C).

The normal membrane structures observed in the *emp24Δ* single mutant cells suggest that vesicle size alterations cannot explain the lax ER retention phenotype (Elrod-Erickson and Kaiser, 1996). Instead, we hypothesized that reduced cargo occupancy in the mutant vesicles might create space that would allow for stochastic capture of ER resident proteins. The CLEM approach does not allow us to visualize cargo proteins within vesicles, so we turned to biochemical analyses to test this hypothesis. We first sought to rule out various indirect effects of a p24 deletion that might result in ER retention defects: (i) chaperone induction via the Unfolded Protein Response (UPR) could saturate ER retrieval (Belden and Barlowe, 2001); (ii) ER resident chaperones could be exported from the ER while bound to misfolded clients, which might be expelled from the ER under stress (Satpute-Krishnan et al., 2014); (iii) increased diffusional mobility of proteins in the ER lumen could promote access to ERES (Lai et al., 2010); and (iv) altered Golgi-ER retrieval could increase secretion.

### ER retention defects in p24 mutant strains are not explained by ER stress-induced effects

We first tested UPR effects by deleting Hac1, the transcription factor responsible for UPR activation, in two p24 mutants, *erp1Δ* and *erp2Δ*, observing robust secretion of the abundant ER lumenal HSP70, Kar2, in a colony immunoblot assay even when the UPR is abrogated (Figure 3A). Since deletion of *EMP24* is inviable in the absence of the UPR (Copic et al., 2009), we sought an independent means to test UPR dependence in the *emp24Δ* strain. Abrogation of the UPR element within the *KAR2* promoter yields a strain, *upre^d^*-*KAR2,* in which *KAR2* expression is uncoupled from ER stress (Hsu et al., 2012). We deleted *EMP24* in this background and found no reduction in Kar2 secretion, confirming that Kar2 leakage in p24 mutant strains is unrelated to UPR induction (Figure 3B). We next tested whether Kar2 secretion in the *emp24Δ* strain results from its ER expulsion in complex with misfolded clients, which might be triggered by ER stress (Satpute-Krishnan et al., 2014). We deleted *EMP24* in a *kar2-1* mutant strain, in which client-binding is abrogated. This strain has a constitutive UPR and hence upregulates Kar2 (Figure 3C, lysate). This condition leads to elevated Kar2 secretion in a wild-type background, but *EMP24* deletion further enhanced Kar2 secretion, suggesting p24-driven release is independent of client interaction (Figure 3C).

**Figure 3.**
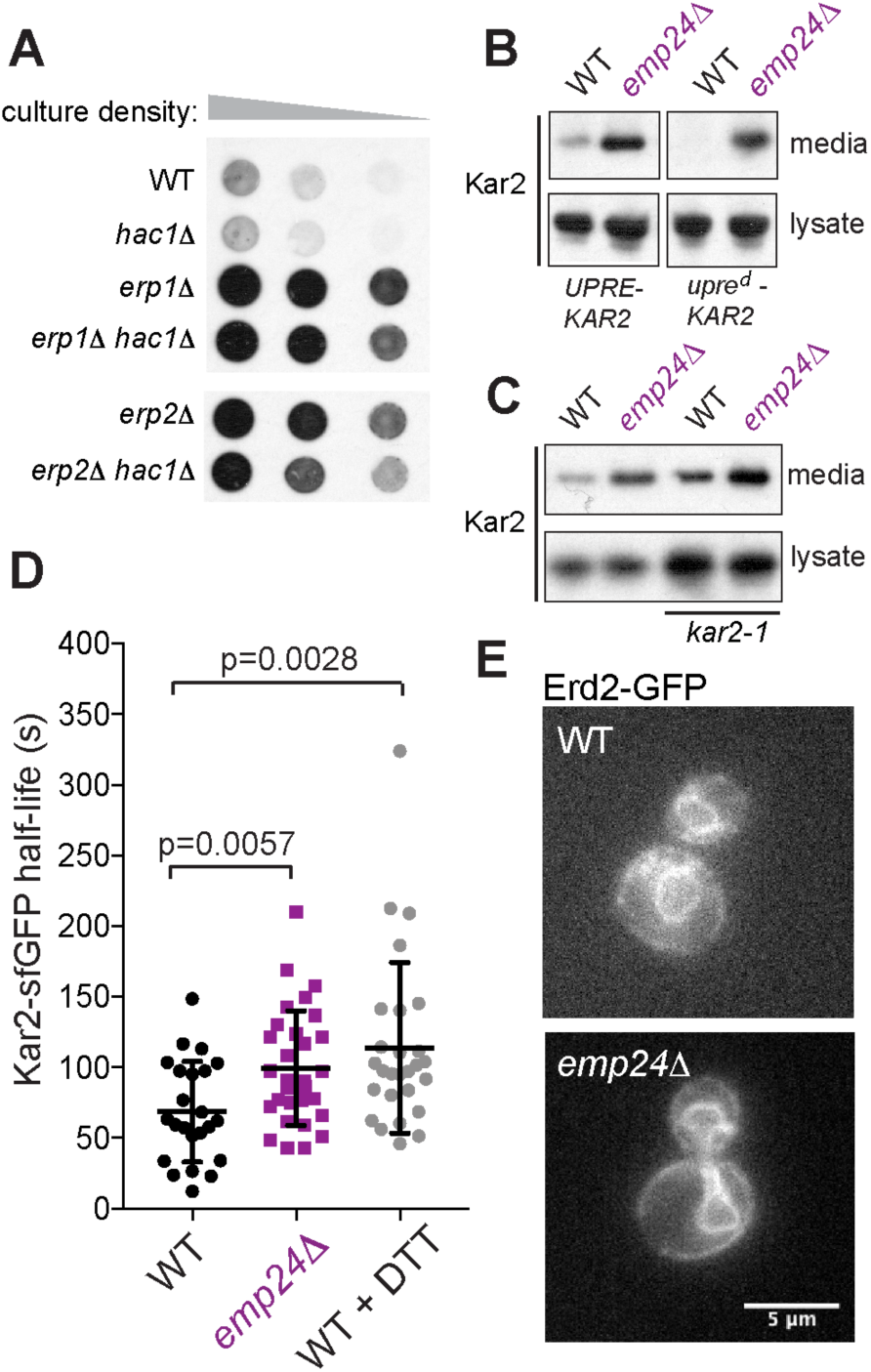
Kar2 secretion is not due to UPR, retrieval failure or changes in ER lumenal mobility. (A) Serial dilutions of the indicated yeast strains were overlaid with nitrocellulose, and secreted Kar2 detected with Kar2-specific antibodies. (B) and (C) Kar2 was detected in intracellular (lysate) and secreted (media) fractions by SDS-PAGE and immunoblotting with anti-Kar2 antibodies. (D) Mobility of Kar2-sfGFP was measured by FLIP. Kar2 half-life in individual cells was measured in the indicated strains. WT + DTT cells were treated with 5 mM DTT for 1 h. The graph shows the mean and the error bars represent SD; n=23 (WT); n=30 (*emp24*Δ); n=27 (WT + DTT). Statistical test was a t-test. (E) Fluorescence microscopy of WT and *emp24*Δ cells expressing Erd2-GFP revealed ER localization in both strains.

To address the diffusional mobility of Kar2 within the ER lumen, we used fluorescence loss in photobleaching (FLIP) to measure the half-life of Kar2-sfGFP. In this assay, a region of the cell containing cortical ER is continuously photobleached, and the loss of fluorescence in regions outside the photobleached region quantified. This gives a measure of the diffusional mobility of Kar2-sfGFP, which has previously been demonstrated to decrease significantly under conditions of ER stress (Lajoie et al., 2012). We similarly observed an increase in Kar2-sfGFP half-life (i.e. decreased mobility) upon treatment with DTT (Figure 3D). The *emp24*Δ mutant also showed reduced Kar2-sfGFP diffusion (Figure 3D), consistent with the known constitutive UPR activation in this strain. This observation that the ER lumen is not more diffusive in *emp24*Δ cells argues against the hypothesis that elevated export of ER resident proteins results from enhanced access to ERES that is normally restrained by limiting the diffusion of these proteins. Finally, we sought to address whether Golgi-ER retrieval of escaped Kar2 by the HDEL-receptor Erd2 was impaired in the *emp24*Δ strain. We examined the localization of Erd2-GFP which normally localizes to the ER due to rapid Golgi-ER traffic (Schuldiner et al., 2005). A similar localization was observed in the p24 mutant *emp24*Δ suggesting that retrieval is functional in these ER-retention mutants (Figure 3E).

### ER-retention mutants show higher rates of bulk flow

Having ruled out various indirect effects of p24 deletion on Kar2 secretion, we aimed to test the model that increased bulk flow could explain leakage of ER residents in these and other mutant strains. We first measured secretion of an inert marker, the C-terminal protease domain, Cp, of the Semliki Forest virus capsid protein. This protein has been previously used as fluid-phase marker as it folds rapidly in a chaperone-independent manner, does not undergo covalent modifications, and is unlikely to possess binding signals for cargo receptors (Thor et al., 2009). We generated a yeast version of this marker that used an Ost1 signal peptide, followed by a FLAG epitope, and the Cp protease domain. We note that the published mammalian construct uses an HA epitope, which reveals a Tyr-Pro-Tyr ER export signal following signal peptide cleavage that might function as a cryptic ER export signal by interaction with SURF4/Erv29 (Yin et al., 2018). Upon galactose induction of the bulk flow marker Cp, we observed significant secretion in wild-type cells, consistent with constitutive bulk flow. In the *emp24*Δ strain, Cp secretion was elevated, suggesting bulk flow is enhanced in this background (Figure 4A). Quantification of Cp secretion using radioactive pulse-chase revealed an increase of ∼24% in the *emp24*Δ strain at t=30 min (Figure 4B). Together, our observations are consistent with increased bulk flow rates as a cause of ER leakage in *emp24*Δ mutants.

**Figure 4.**
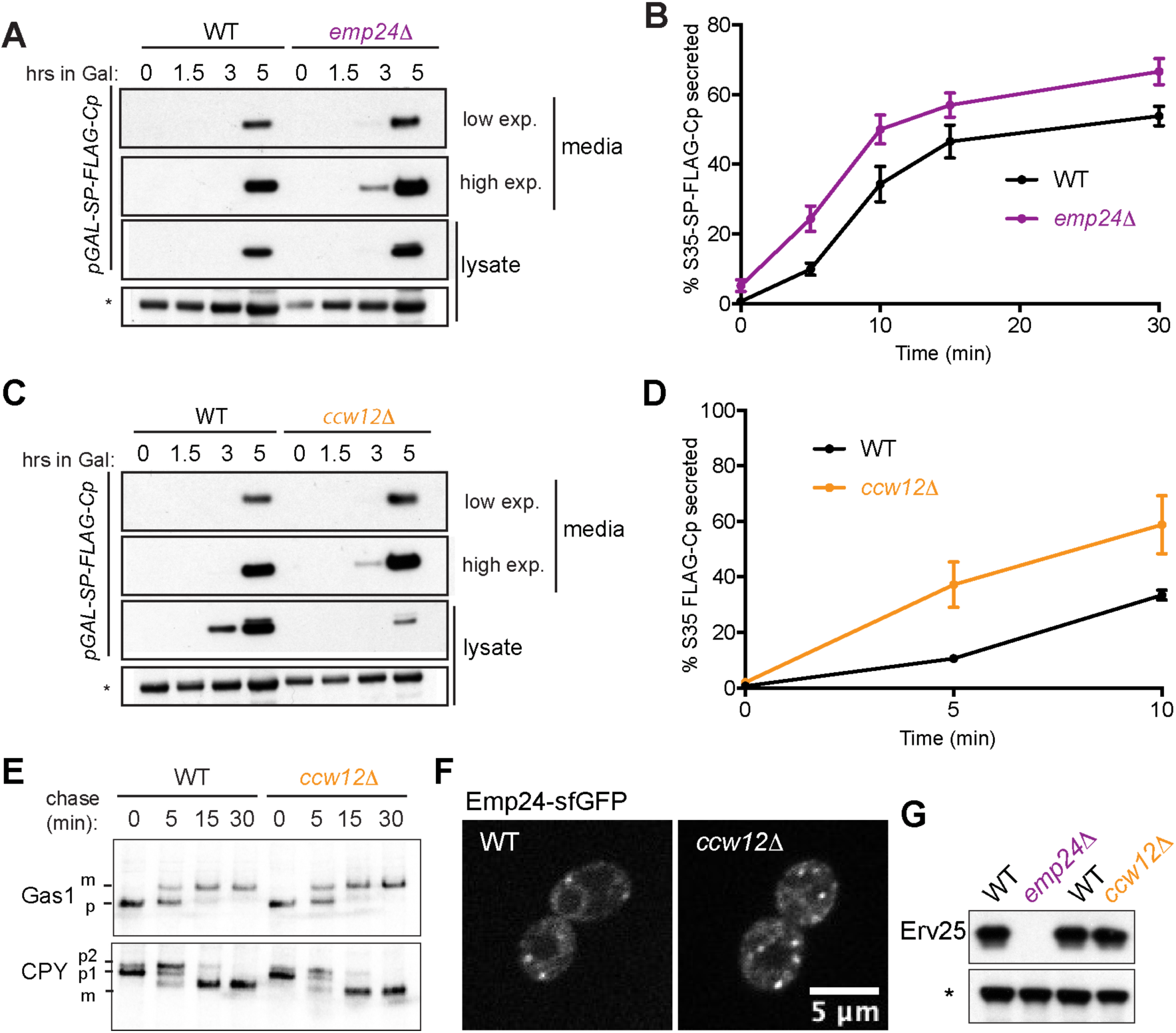
Elevated bulk flow stems from decreased cargo crowding. (A) WT and *emp24*Δ strains expressing *GAL_pr_-SP-FLAG-Cp* were induced with 0.02% galactose, and separated into intracellular (lysate) and extracellular (media) fractions. Cp was detected by SDS-PAGE and anti-FLAG immunoblot. (B) FLAG-Cp was immunoprecipitated from media and lysate fractions of [^35^S]methionine-labelled cells at the indicated times, analyzed by SDS-PAGE and detected by autoradiography. The percentage of secreted SP-FLAG-Cp is plotted for the indicated times. Error bars depict SEM; n=6. (C) as described in A. (D) as described in B. Error bars depict SEM; n=3. (E) Gas1 and CPY maturation were examined in wild-type and *ccw12Δ* strains by pulse-chase with [^35^S]methionine. Gas1 and CPY were immunoprecipitated from lysates at the indicated times and detected by SDS-PAGE and autoradiography. (F) Fluorescence microscopy of WT and *ccw12Δ* cells expressing Emp24-sfGFP revealed similar localization of Emp24. (G) Steady state levels of Erv25 in the indicated strains were measured from whole cell lysates by immunoblotting using an Erv25-specific antibody. A non-specific band labelled with an asterisk is shown as loading control.

We next sought to test whether enhanced bulk flow was a unique feature of the p24 mutants, or whether this model could also apply to ER retention mutants that are not integral parts of the ER-Golgi trafficking machinery. Ccw12 is a highly abundant cell wall GPI-AP, comprising ∼12% of the GFP-secretome (Ghaemmaghami et al., 2003; Caro et al., 1997) (Figure S3A). Deletion of *CCW12* causes Kar2 secretion (Figure S3B and (Copic et al., 2009)), so we reasoned that the simple loss of this one cargo protein may mimic the p24 mutant condition by creating empty space in a vesicle to permit non-specific capture. Indeed, secretion of FLAG-Cp was increased in a *ccw12Δ* strain (Figure 4C and 4D), consistent with increased bulk flow leakage of ER lumenal proteins. Loss of Ccw12 had no impact on Kar2 mobility (Figure S3C) or Erd2 localization (Figure S3D), and ER export rates of the vacuolar protease, CPY, and the cell wall glycoprotein, Gas1, were normal (Figure 4E). Normal Gas1 secretion is indicative that p24 function itself is not compromised in the *ccw12Δ* strain, but we further excluded that ER leakage might be indirectly caused by aberrant p24 function by examining recruitment of Emp24-sfGFP to ERESs, which was normal (Figure 4F). Moreover, deletion of *CCW12*, unlike *EMP24* deletion, did not destabilise the other main p24 protein Erv25 (Figure 4G). Together, the phenotypes of the *ccw12Δ* strain suggest that simple cargo occupancy can explain the decreased stringency of ER export sorting in p24 and other mutants. We propose that enrichment of abundant cargo proteins, especially glycosylated cell wall proteins that are likely to occupy significant space, creates steric pressure within the vesicle lumen that excludes ER resident proteins and diminishes bulk flow.

### Vesicle occupancy restrains bulk flow

If Kar2 secretion is a consequence of increased bulk flow caused by increased lumenal volume available for stochastic capture, then modulating cargo occupancy should influence bulk flow. In this context, we sought to test whether p24 proteins have a specific role in modulating sorting stringency at the ER or whether local concentration of cargo directly restrains bulk flow. We tested the effects of specific p24 domains on bulk flow by using chimeric proteins that comprised a cleavable signal peptide and various substitutions of lumenal and transmembrane domains of Emp24 (Figure 5A). All constructs preserved the short cytosolic ER domain that contains the export signal that drives incorporation into a vesicle. We replaced the lumenal GOLD (for <u>Gol</u>gi <u>d</u>ynamics) domain with sfGFP, generating the chimeric protein GFP-CC-TM. We also deleted the short helical coiled-coil region, thought to participate in p24 oligomerization (GFP-TM), and replaced the transmembrane domain with a generic 26 Leu repeats (GFP-26xLeu). Each of these constructs was introduced into an *emp24*Δ strain, driven by the strong *GAL1* promoter. All proteins could be visualized in the ER, plasma membrane, and vacuole (Figure S4A)consistent with ER export and onward traffic. We monitored Kar2 secretion upon galactose induction, observing a decrease in extracellular Kar2 as the levels of the chimera proteins, serving as cargo, increased (Figure 5B). None of the chimeras stabilized Erv25, suggesting they are not capable of functional oligomerization (Figure S4B). That each chimera was capable of restoring sorting stringency suggests that simple cargo occupancy in a vesicle, rather than specific functions of the p24 proteins, is what drives selectivity. Whether organization into an array-like structure upon coat binding also contributes to this selectivity filter remains to be further explored. We note that enrichment of these chimeric cargo proteins in vesicles is essential for their reversal of ER leakage. Similar reversal was not observed when other soluble bulk flow cargoes were tested for competition (Figure S4C).

**Figure 5.**
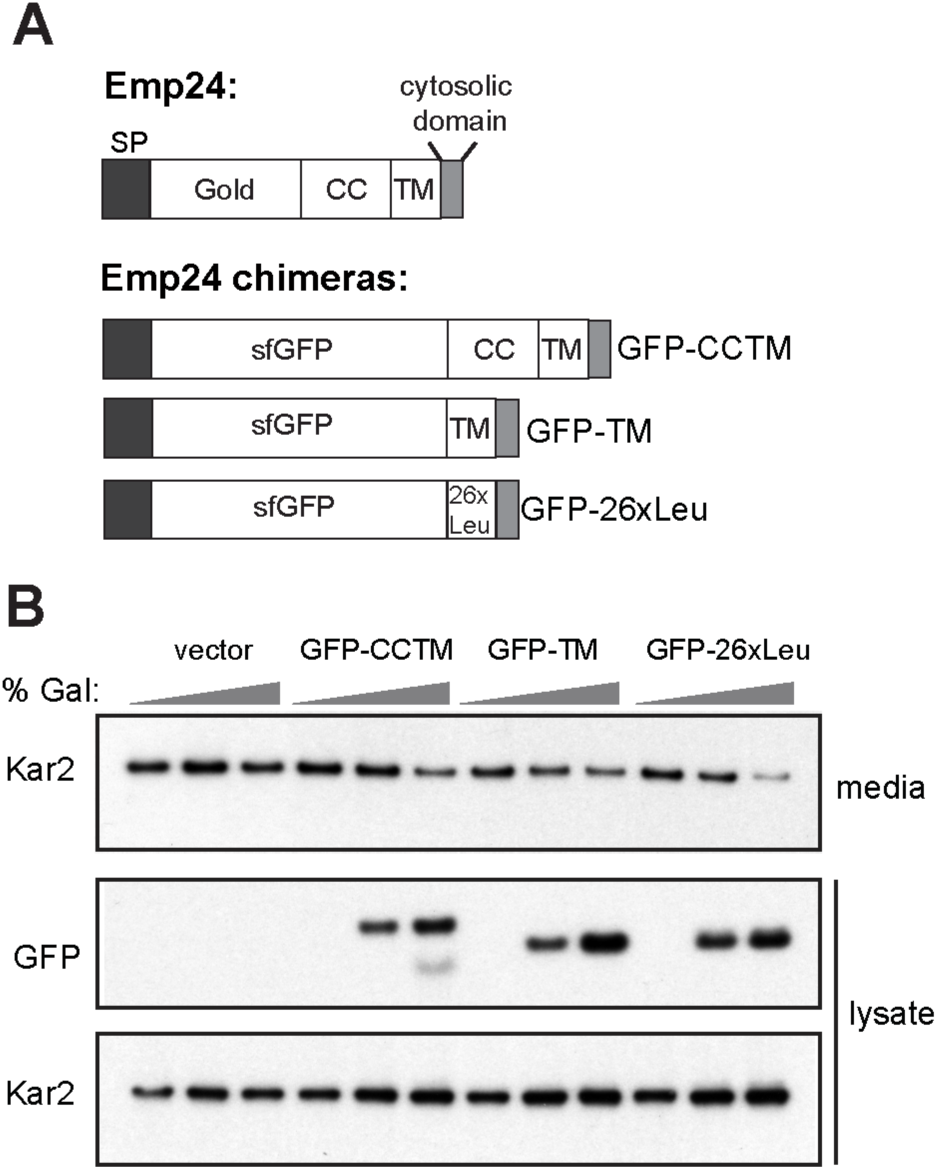
Restoring cargo occupancy reverses ER leakage. (A) Schematic of Emp24 and chimeras used in B. (B) Kar2 secretion was analysed in *emp24*Δ cells after galactose induction of the constructs indicated. Intracellular (lysate) and extracellular (media) proteins were resolved by SDS-PAGE and detected by Western blot against Kar2 and GFP.

### Bulk flow is modulated by vesicle size

If cargo occupancy is a main constraint on bulk flow, then vesicle size should also influence efficiency of stochastic cargo capture. Specifically, reducing vesicle size should restore a steric constraint. In yeast, COPII vesicle size is influenced by the cargo adaptor layer; vesicles formed *in vitro* with the Sec24 paralog Lst1/Sfb3 are ∼15% larger in diameter than those formed with Sec24 (Miller et al., 2002; Shimoni et al., 2000; Lee et al., 2002). Thus, *lst1Δ* cells should generate small Sec24-only vesicles thereby imposing a greater steric effect on the vesicle lumen. We examined Kar2 secretion in *emp24*Δ and *ccw12Δ* strains in the absence and presence of Lst1. Deletion of *LST1* reversed Kar2 secretion in these backgrounds, consistent with restoration of steric pressure and subsequent decrease in bulk flow leakage. This reversal was rescued by reintroduction of either *LST1* or *lst1-B*, which is mutated in the cargo-binding site (Figure 6A). This suggests that the effect of *LST1* deletion is related to its structural role rather than cargo capture.

**Figure 6.**
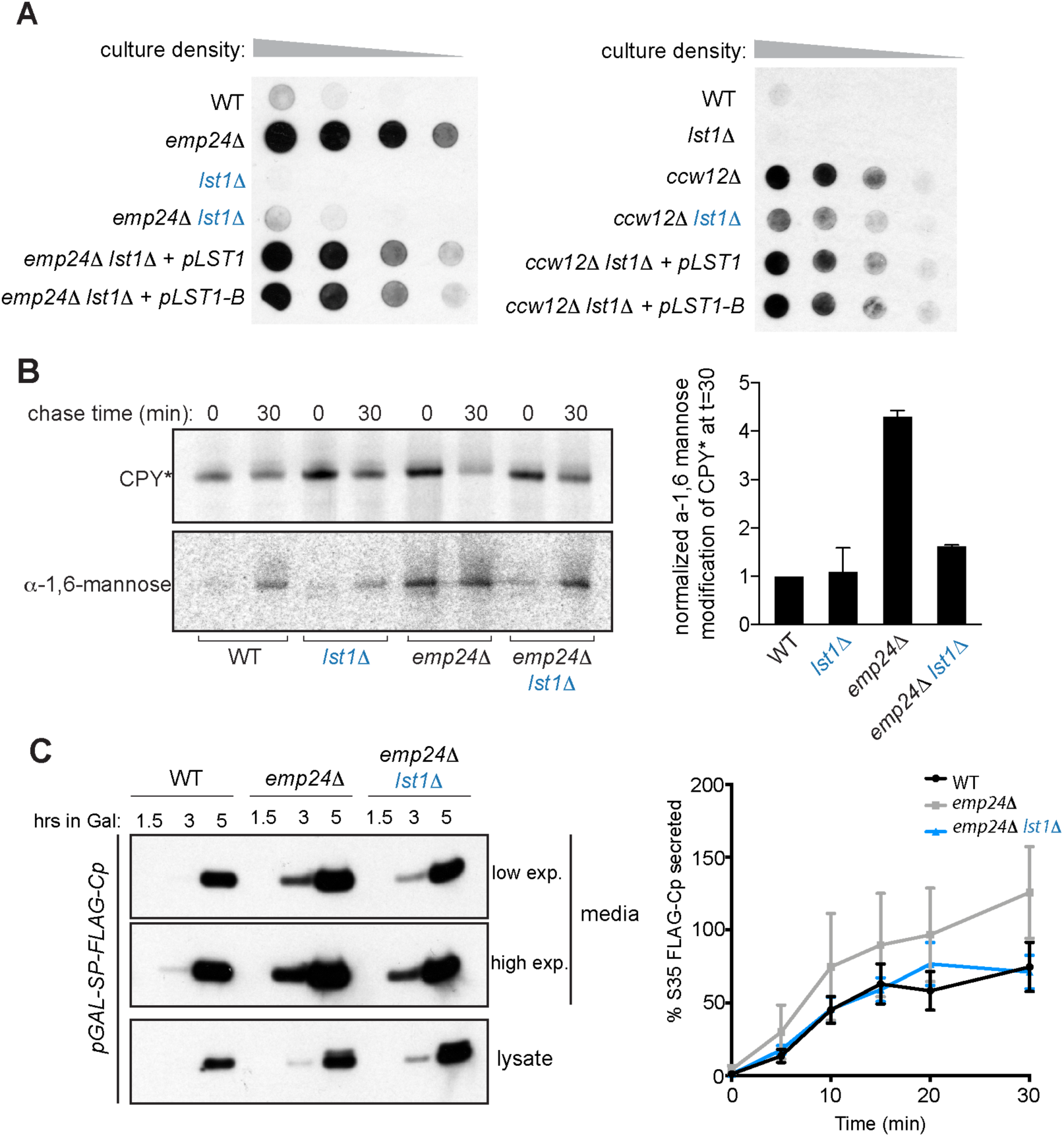
*LST1* deletion restores cargo stringency and reduces bulk flow. (A) Serial dilutions of the indicated strains were spotted onto YPD plates and Kar2 secretion determined by colony immunoblot as described in Figure 3A. (B) Wild-type, *lst1Δ, emp24*Δ and *emp24Δ lst1Δ* cells expressing HA-CPY* were subjected to pulse-chase analysis. CPY* was immunoprecipitated from lysates at the indicated times and either analysed directly or subjected to secondary immunoprecipitation using anti-*α*-1,6-mannose antibodies. Immunoprecipitated proteins were resolved by SDS-PAGE and detected by autoradiography. The ratio of Golgi-modified CPY* to total CPY* relative to a wild type strain at t=30 min was determined for three independent experiments. Averages and standard deviations are plotted. (C) FLAG-Cp was detected in strains indicated after induction with 0.02% Galactose as described in Figure 4A (left panel). Pulse-labeled proteins were immunoprecipitaded from the media and lysates at the indicated times as described in Figure 4B. Error bars depict SEM; n=3 (right panel).

Since steady-state Kar2 secretion is influenced by both anterograde and retrograde pathways, we sought to test whether the *lst1Δ* effect was specific for COPII-mediated forward traffic. We thus examined leakage of misfolded CPY*, which is another phenotype of the *emp24*Δ condition (Copic et al., 2009). The degree of CPY* trafficking to the Golgi is measured by acquisition of an *α*-1,6-mannose sugar, which represents a quantitative readout of delivery to the Golgi lumen. In wild-type and *lst1Δ* cells the misfolded protein fails to reach the Golgi and is rapidly degraded, resulting in low levels of *α*-1,6-mannose modification. In *emp24*Δ cells, *α*-1,6-mannose-modified CPY* was increased, as previously reported, whereas deletion of *LST1* reversed this effect (Figure 6B). Finally, we confirmed that deletion of Lst1 impacts the anterograde pathway by measuring bulk flow of SP-FLAG-Cp. Compared with the elevated levels of Cp secretion in *emp24*Δ cells, the *emp24Δ lst1Δ* double mutant showed bulk flow rates identical to wild-type cells (an average reduction of 18% relative to the *emp24*Δ single mutant) (Figure 6C).

Having demonstrated that deletion of *LST1* indeed restores sorting stringency, we sought to quantify the effect of loss of Lst1 on vesicle size *in situ* using CLEM. We acquired 37 tomograms at sites of COPII vesicle formation in a *lst1*Δ *emp24*Δ *SEC16-sfGFP* strain (Figure 7A). Most of these ERES had more than one adjacent free vesicle, which we segmented to quantify for volume. Vesicles in the *lst1*Δ *emp24*Δ strain had a median volume of 31,493 nm^3^ (Figure 7B), corresponding to a volume reduction of 21% compared to *emp24*Δ. The majority of COPII vesicles in the *lst1*Δ *emp24*Δ strain fell below the median vesicle volume of wild type *SEC16-sfGFP* and *emp24*Δ *SEC16-sfGFP* cells. The smallest vesicles for all strains tested had similar volumes, which highlights that the formation of large COPII vesicles is specifically impaired when Lst1 is deleted.

**Figure 7.**
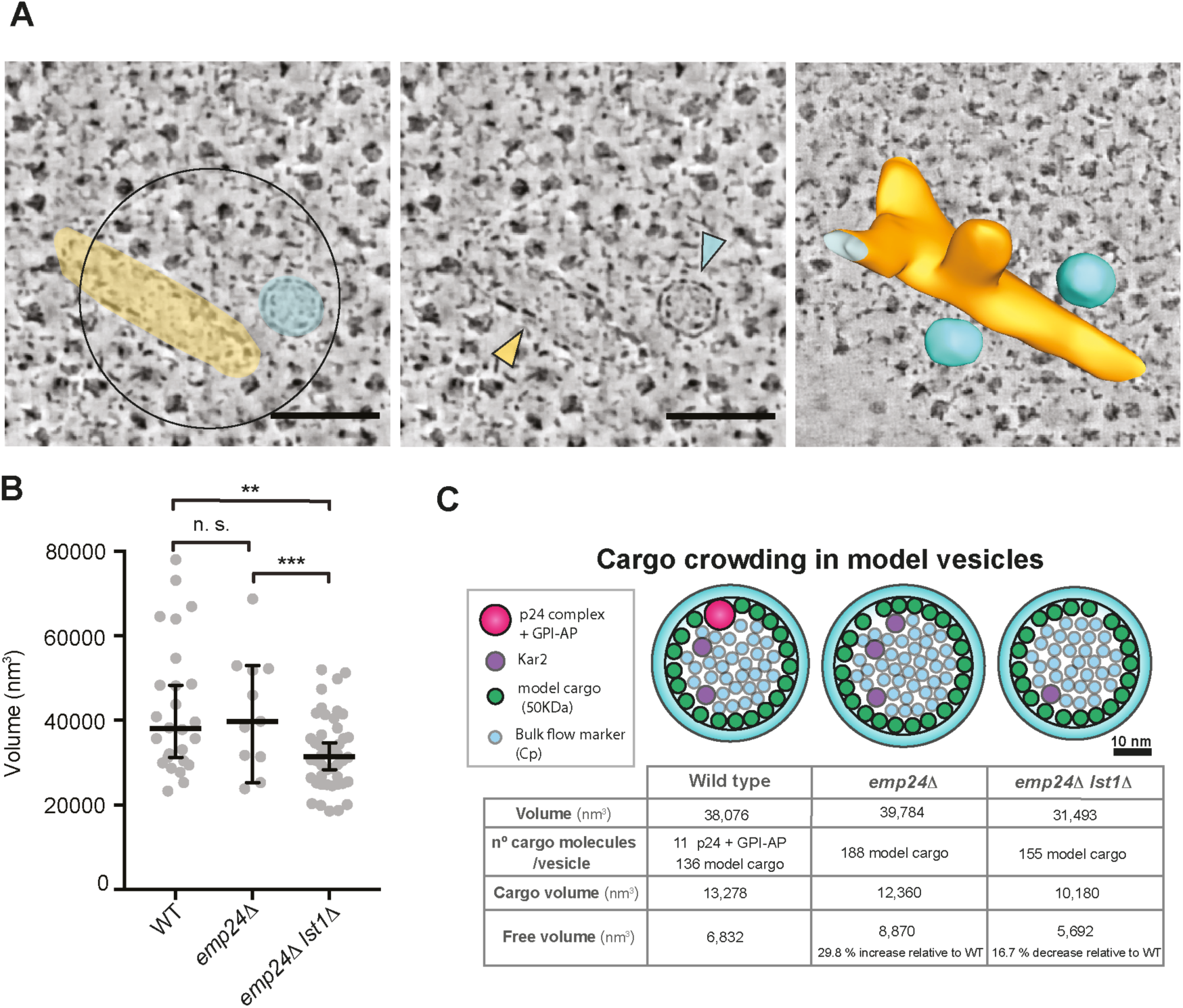
The volume of COPII vesicles is significantly reduced in *emp24*Δ *lst1*Δ cells. (A) A Sec16-sfGFP positive ERES in an *emp24Δ lst1Δ* cell. Left panel is a virtual tomographic slice showing a false-coloured ER tube (yellow) and a free vesicle (cyan). Central panel shows the same structures highlighted by coloured arrowheads. Right panel is a segmentation model of the corresponding 3D membrane ultrastructure showing a bud and two vesicles. Scale bar is 100 nm. (B) Plot of the volume (nm^3^) of COPII vesicles in the strains indicated. Bars correspond to median value and 95% confidence interval. Statistical test was a t-test, ** p<0.001, *** p<0.0001. (C) Model of cargo crowding in vesicles. 2D sections of vesicles drawn to scale illustrating the cargo crowding effects in COPII vesicles. The vesicle volume and volume of cargo in a given vesicle determines the free volume susceptible to bulk flow. Accordingly, p24-depleted vesicles (*emp24*Δ) or smaller size vesicles (*emp24Δ lst1Δ*) differ in the free space available (see methods for calculation details).

Together our data suggest that cargo occupancy in ER-derived vesicles inversely correlates with the degree of bulk flow, and that the effect of cargo occupancy is to create steric effects that may serve in part to prevent inappropriate export of ER residents. Since our 3D measurements of vesicle volume in cells permits a quantitative analysis, we sought to model the effects of vesicle size and cargo occupancy on ER sorting (Figure 7C). We reasoned that actively recruited cargo proteins, engaged with Sec24/Lst1, will occupy the volume immediately underlying the membrane, and that the surface area of this region sets a limit on the number of cargoes that can be loaded. For simplicity, we considered only two classes of cargo proteins, each modelled as spheres: the p24 complex consisting of four p24 proteins together with the GPI-AP, Gas1, and a model “average” cargo of 50 kDa (Lodish et al., 2012). Since GPI-APs make up ∼25% of the GFP-secretome (Figure S3A), we assigned one-quarter of the vesicle volume to the p24/GPI-AP complex, with the remaining inner membrane surface area occupied by the model cargo. The cargo-occupied volume was subtracted from the total lumenal volume to yield the “free volume” that is accessible to non-selective capture. Comparing the wild-type condition with that of an *emp24*Δ strain, in which the p24/GPI-AP volume is absent, we calculate an increase of ∼80% in free lumenal volume. If we assume that absence of the p24/GPI-AP complex results in free membrane surface that can be occupied by additional “model” cargo proteins, then the increase in free volume is ∼30%, a value similar to the experimentally determined ∼24% increase in Cp secretion (Figure 4B and Figure 6C). In the *emp24Δ lst1Δ* double mutant, the reduction in vesicle size translates to a reduction in free volume of ∼17% compared to wild type and ∼36% relative to *emp24*Δ cells. This is a larger reduction than we observed for Cp secretion, which revealed equivalent rates of bulk flow in the *emp24Δ lst1Δ* and wild-type strains (Figure 6C). Our model does not account for additional factors that are almost certain to influence the behaviour of individual proteins with respect to their ability to access the bulk fluid, including folding state, glycosylation, and hydration volumes. Indeed, the different effects of *LST1* deletion on Cp secretion and Kar2 leakage serve to highlight such distinct behaviours: Kar2 secretion was completely reversed in the *lst1Δ* background, whereas the effect on Cp secretion was more modest. Nonetheless, this simplified model provides support for our proposal that cargo loading and vesicle size contribute to steric effects that influence ER export.

## Discussion

ER export of nascent secretory proteins represents an important quality control checkpoint, whereby immature and misfolded proteins, along with ER resident proteins, can be excluded from vesicles as a mechanism of ER retention. Emerging evidence suggests that cells have evolved complex signalling pathways to modulate vesicle formation in the context of cargo burden (Subramanian et al., 2019; Centonze et al., 2019). Such adaptation is probably particularly important to address cell-specific needs, and physiological responses to changing environments. Here, we describe an additional stochastic mechanism of regulating ER export, which derives from the principle that secretory cargo are not inert passengers, but can confer biophysical constraints on the vesicle formation process. As a result, the steric pressure derived from molecular crowding prevents improper capture of non-specific cargo.

### Vesicle morphology and cargo occupancy as drivers of quality control

Our visualization of membrane morphology by CLEM revealed that in the absence of bulky cargo proteins and Sec13, ERES became more pleiomorphic, with individual vesicles generally larger in size. However, the lumenal volume of p24-depleted vesicles when Sec13 was present was similar to that of wild-type vesicles, suggesting that the observed increase in bulk flow upon p24 depletion alone is driven by a reduction of cargo occupancy in these vesicles rather than a larger vesicle lumen. As abundant constituents of COPII vesicles, the p24 proteins (and associated cargoes) may act simply as molecular ballast to occupy space and thus exclude ER residents and minimize bulk flow. Furthermore, the capacity of these proteins to oligomerize, in coordination with the coat, could additionally facilitate the exclusion of non-cargo proteins through formation of a diffusion barrier (Ma et al., 2017). We find that reducing vesicle size in the p24-depleted background restores the available volume to wild-type levels, and leads to decreased bulk flow even in the absence of p24 function (Figure 6). Additional support for this simple steric model comes from our observation that sorting stringency was also restored by an artificial increase in cargo crowding (Figure 5). The fact that deletion of another abundant cargo, the cell wall mannoprotein Ccw12, similarly caused ER leakage indicates that enforcing sorting stringency is not an exclusive property of p24 proteins, but can be imposed by the abundance of other bulky cargo. We thus propose that ER retention of residents and minimization of bulk flow is a more general effect arising from the limited available volume inside a budding vesicle and active cargo selection by the coat.

### Coat composition drives vesicle morphology

In order to minimize ER leakage, a cell could reduce the number and size of vesicles to ensure maximal cargo packing. However, some large cargoes have spatial needs that might not be accommodated by small vesicles, necessitating a larger carrier. Large vesicles risk excessive bulk flow if the cargo capacity is not met, thus cargo-driven modulation of vesicle size via coat adaptors represents a simple model by which secretion can be tuned to reflect cargo needs. Large cargoes, such as the plasma membrane H^+^-ATPase, Pma1, as well as GPI-APs, have a preference for Lst1 (Roberg et al., 1999), which generates bigger vesicles than Sec24 alone. Thus, cross talk between cargo and coat might tailor vesicle size to fit cargo exigency while still maintaining high sorting stringency. The mechanism by which different cargo adaptor isoforms yield vesicles of distinct size remains to be seen, but our CLEM analysis confirms that in cells, depletion of Lst1 indeed results in a significant reduction of vesicle volume. Given the capacity for the inner COPII coat to form oligomeric arrays, it is possible that Lst1 contributes directly to curvature generation to both form a larger structure and provide force to counter the physical effects of its large clients.

Similar cargo-driven programming of vesicle size is likely to occur in mammalian cells. The collagen export receptor, TANGO1, has been proposed to drive coat organization to favour tubule formation around an export-competent collagen fibre. However, other modes of collagen export have also been proposed, where local membrane fusion and remodelling might occur (Malhotra and Erlmann, 2015; McCaughey et al., 2019). Whether and how ER resident proteins are excluded from such structures remains to be fully explored. Indeed, in some cases, minimization of ER leakage may not be important; in professional secretory cells, abundant cargo may be packaged at the prevailing ER lumenal concentration. If lack of cargo enrichment leads to ER leakage, retrieval pathways might compensate, or the cell might simply tolerate a degree of loss of ER residents.

In addition to the inner coat adaptors, the outer coat scaffold also contributes force to bend a cargo-replete membrane. Traffic of large cargo is more dependent on Sec13 (Townley et al., 2008), and reduced packaging of such cargo is permissive to loss of Sec13 (Copic et al., 2012). Our CLEM analysis in such cargo-depleted yeast cells with a Sec31-only cage revealed an increase in vesicle size, consistent with a reduced ability of Sec31 to enforce curvature. However, smaller vesicles of 45 to 55 nm in diameter were still observed in the absence of Sec13, suggesting Sec31 alone can achieve high curvatures. Whether this smaller population of vesicles has a distinct cargo composition that is more permissive to curvature generation remains to be determined. Another feature of ERES under Sec31-only conditions was the increased number of buds at Sec16-correlated sites. This may be a consequence of frustrated budding events where Sec31 fails to deform the membrane to a fission point. If this is the case, it suggests a second structural role for Sec13 in ensuring adequate curvature to drive timely vesicle fission.

Our model of cargo-driven steric effects driving ER sorting stringency has interesting implications for the secretory pathway more broadly, as well as for physiological regulation of protein traffic. To preserve ER quality control, cells may regulate vesicle formation and vesicle size to match cargo burden. Specific signalling cascades that modulate vesicle formation according to cargo requirements have been recently described (Subramanian et al., 2019; Centonze et al., 2019), and such pathways may tune ERES size and abundance in the context of cargo abundance. Whether similar principles act at other trafficking routes remains to be tested. Fluid phase uptake from the cell surface is well-established but may differ under conditions where active cargo capture is enhanced, or vesicle formation becomes limiting. Biophysical effects in clathrin-independent endocytosis are of particular interest, where uptake of proteins can be determined by steric bulk (Bhagatji et al., 2009). Diversion of such cargo to non-clathrin pathways may permit bulk flow uptake via clathrin-dependent mechanisms. Dissecting these different pathways, and exploring steric effects in vesicles more broadly should shed further light on how cells manage the stringency of protein delivery.

## Materials and Methods

### Strains and plasmids

All strains were constructed and grown using standard *Saccharomyces cerevisiae* methods. Strains (Table 1) were made by crossing, sporulation, and dissection of tetrads or by PCR-based integration of auxotrophic or drug-resistance markers. Plasmids used in this study are listed in Table 2. The *pSP-GFP-TM* construct containing the Kar2 signal sequence followed by superfolder GFP and the transmembrane and cytosolic domains of Emp24 (residues 153-203) was purchased as a synthetic construct in pRS316 (*GeneScript*). The *pSP-GFP-CCTM* plasmid contains the same sequence with the addition of the coiled-coil sequence of Emp24 (residues 129-152) upstream of the transmembrane domain, and was also synthesized commercially (*GeneScript*). The *pSP-GFP-26xLeu* construct was created from *pSP-GFP-TM*, where the Emp24 transmembrane domain (residues 173-193) was replaced by 26 leucines using Gibson assembly (*New England Biolabs*). To construct *pSP-FLAG-Cp,* the sequence containing the HA epitope and that of the Semliki forest virus capsid protease domain (Cp) was amplified from plasmid p626 (a gift from Ari Helenius) and ligated into pRS426. In the resulting plasmid, pRS426-HA-Cp, the HA sequence was replaced by the Ost1 signal peptide followed by the FLAG epitope, using sequential reactions of QuikChange multi site-directed mutagenesis (*Agilent*).

### Correlative Light Electron Microscopy (CLEM)

CLEM was performed as described in (Kukulski et al., 2011) with modifications described in (Ader and Kukulski, 2017). Yeast cells were grown at 25° C in SC-TRP medium to 0.6-0.8 OD_600_ and pelleted by vacuum filtration on nitro-cellulose discs, then placed on an agar plate to prevent the pellet from drying out. The resulting yeast paste was high-pressure frozen in 200 µm deep wells of aluminium carriers (*Wohlwend*) using a HPM100 (*Leica Microsystems*). Freeze-substitution and Lowicryl HM20 (*Polysciences, Inc.*) resin-embedding were done as previously described in (Kukulski et al., 2011). 0.03% uranyl acetate in acetone was used for freeze-substitution. Samples were shaken on dry ice for the first 2-3 h of freeze-substitution. Sections of 300 nm thickness were cut with an Ultra 45° diamond knife (*Diatome*) on an Ultracut E microtome (*Reichert*). The sections were floated onto PBS and picked up with 200 mesh carbon-coated copper grids (*AGS160, Agar Scientific*). Fluorescent TetraSpeck beads (*Invitrogen*), 50 nm in diameter, were adsorbed onto the grid. Directly after sectioning, grids were mounted for fluorescence microscopy (described below). Prior to electron tomography, 15 nm gold beads (*Electron Microscopy Sciences*) were adsorbed on the sections, which were then post-stained for 15 min with lead citrate. Scanning transmission EM tomography was done on a TF20 microscope (FEI) with an axial brightfield detector, using a camera length of 200 mm and a 50 mm C2 aperture (Ader and Kukulski, 2017; (Hohmann-Marriott et al., 2009). For correlation to fluorescence images, low magnification tilt series at 3.1 nm pixel size were acquired using SerialEM (+/-55° tilt range, 2° increment, single axis acquisition) (Mastronarde, 2005). Higher magnification tomograms were acquired with dual axis tilt series +/-60° with 1° increment and at 1.1 nm pixel size (Mastronarde, 1997). All tomographic reconstructions were done in IMOD (Kremer et al., 1996) and fiducial-based correlation was done using MATLAB-based scripts described in (Kukulski et al., 2011).

### Segmentation analysis

The 3D membrane morphologies of correlated ERES shown in the Figure panels were segmented by manual tracing, simplification and smoothening of the resulting surfaces using Amira (*Thermo Fisher*). Thereby generated models are for illustration purpose only.

Vesicles diameter and volume quantifications were obtained with FIJI using the plugin LimeSeg (Machado et al., 2019) as following: the outer contour of a vesicle was selected to the ROI using the “point tool” and “segmented line” tool moving in z through the tomographic slices, adding to the ROI the lowest plane of the vesicle with the point tool, then clicking contours with the segmented line tool every 5 virtual slices and finally closing the volume selecting the top plane of the vesicle with the point tool. LimeSeg Skeleton Segmentation tool settings were adjusted to recognize and segment the outer surface of the vesicle (D_0: 4, F_pressure: 0, Z_scale: 1, Range_in_DO_units: 1, NumberOfIntegrationStep: -1, RealXYPixelSize: 1). After running the segmentation, the correct distribution of surfels over the outer contour of the vesicle was assessed by eye. The LimeSeg segmentation tool provides the list of vertices of the mesh. The centroid of this point cloud gives an estimate the centre of the segmented vesicle. The maximum radius is then computed as the maximum distances to this centre taking into account the 1.1 × 1.1 × 1.1nm voxel size.

### Secretion assays

To examine Kar2 secretion on plates serial dilutions of logarithmic phase cultures were spotted onto YPD plates and incubated at 30°C for 5 h, at which point colonies were overlaid with nitrocellulose filters and incubated for a further 1 h. Nitrocellulose filters were washed, blocked and incubated with *α*-Kar2 polyclonal sera (provided by Randy Schekman, UC Berkeley). Secreted Kar2 was detected with HRP-conjugated goat-anti-rabbit antibodies followed by ECL detection (*Pierce, Rockford, IL*). Where protein secretion was monitored from liquid cultures, stationary phase cultures were back diluted into fresh media and grown for 5h. 2 OD_600_ units of logarithmic phase cells were collected by centrifugation at 14,000 *g* for 5 min and 1.5 ml of the supernatant fluid collected. Proteins in the media fraction were precipitated by adding 0.15 ml of 100% trichloroacetic acid (TCA) (*Sigma Chemical, St Louis, MO*) and incubated on ice for 30 min. Precipitated proteins were collected by centrifugation, washed with acetone, dried, and resuspended in 30 µl of SDS-PAGE sample buffer supplemented with 50 mM Tris pH 9.4. One-fifth of the sample was resolved by SDS-PAGE for Kar2 immunoblots and one-third for SP-FLAG-Cp or SP-GFP immunoblots. Cell pellets from the 2 OD_600_ units were lysed to obtain whole cell preparations. One-tenth was analyzed by immunoblot.

### FLIP assays

FLIP was used to measure mobility of Kar2-sfGFP. Imaging was performed in cells grown to mid-log phase at 30°C in minimal media lacking tryptophan. Images were taken on an Andor Revolution Spinning Disk microscope with a 40x/1.3NA oil immersion objective and an EMCCD camera. Images were collected using the Andor iQ3 software. A small region of interest was repeatedly photobleached, and the fluorescence intensity of the whole cell was measured for a loss of signal, representing protein that had diffused into the bleaching area. Fluorescence intensity was measured using Fiji and statistical analysis was performed with Prism 7.0 (*GraphPad Software*).

### GFP imaging

Emp24-GFP and Erd2-GFP imaging was performed in cells grown to mid-log phase at 30°C in minimal media lacking tryptophan. Images were taken on an Andor Revolution Spinning Disk microscope with a 40x/1.3NA oil immersion objective and an EMCCD camera. For imaging of Sec16-sfGFP and Sec24-sfGFP, cells were grown at 25°C in minimal media lacking tryptophan. Images were taken on a Nikon Ti2 with a 100x/1.49NA Oil (TIRF) objective and an sCMOS camera. The same imaging methodology was used for the imaging of EM grids with section of resin-embedded cells.

### Western blot

Total protein extracts prepared by alkaline lysis of exponentially growing yeast were separated by SDS-PAGE. Proteins were detected with corresponding antibody and chemiluminescence was visualized according to the manufacturer’s instructions (*ECL Advanced; GE Healthcare*).

### Pulse chase analysis of protein secretion and trafficking

Pulse-chase experiments were used to monitor secretion of the bulk flow marker SP-FLAG-Cp and intracellular transport of Gas1, CPY and the misfolded form of CPY, CPY*. These experiments were performed as previously described (Pagant et al., 2007). Briefly, strains were grown to mid-log phase at 30°C, starved for 15 min, and labeled for 5 min with with 1 μL per OD of cells of EXPRESS ^35^S Protein Labeling Mix (*PerkinElmer*) for 5 or 10 min. The label was chased with excess rich media and 2 OD aliquots of cells harvested at different times. Cells were lysed in detergent and the protein of interest was immunoprecipitated from cell lysates, and cell media (when measuring SP-FLAG-Cp secretion) separated by SDS-PAGE, and detected by phosphorimaging using a Typhoon scanner (*GE Healthcare*). The protein bands were quantified using Fiji and the percentage of the mature or secreted band in each sample was plotted with Prism 7.0 (*GraphPad Software*).

### Vesicle volume model calculations

Our goal was to estimate the cargo occupancy of a vesicle and thereby determine how cargo enrichment and vesicle size might influence bulk flow. Using our quantitative segmentation analysis, we obtained the median vesicle volume for wild-type, *emp24*Δ and *emp24*Δ *lst1*Δ strains. For simplicity, we assumed these segmented volumes to be spheres. To obtain the values of inner vesicle volume and inner vesicle surface, we calculated the radius of the vesicle from the median volume and subtracted 4 nm corresponding to the lipid bilayer. With this new radius, we back-calculated the volume of the corresponding sphere, obtaining the inner vesicle volume and surface area of the lumenal leaflet. To calculate the number of molecules that could occupy the inner vesicle surface, we calculated the maximal surface area occupied by model cargoes. We approximated the volume of model cargoes from their molecular weight using the online tool http://www.calctool.org/CALC/prof/bio/protein_size. The molecular weight of a cargo-engaged p24 complex was estimated as 4 × 25 kDa + 100 kDa (for Gas1 as determined by SDS-PAGE); as an average model cargo we used 50 kDa (Lodish et al., 2012). From the radius of the modelled protein spheres, we calculated the surface area of the different cargoes. The proportion of p24+Gas1 versus other cargo was obtained from the GFP fluorescence of N-terminally tagged secretome proteins (Figure S3A). We divided the inner vesicle surface by the corresponding cargo-surface share, obtaining a number of p24+Gas1 molecules and model average protein. Knowing the number of molecules that populate the inner surface of the vesicle, we multiplied the number of these proteins by their respective estimated volume to obtain the cargo-occupied volume. Subtracting the volume occupied by cargo from total lumenal vesicle volume, we obtained the available lumenal volume or free space inside the vesicle.

## Supplemental material

Two supplemental tables describe the yeast strains and plasmids used in this study.

## Acknowledgments

We thank Randy Schekman, Davis Ng, Ari Helenius, Jeff Brodsky and Erik Snapp for providing strains, plasmids and antibodies. We thank the LMB light microscopy and electron microscopy facilities for their assistance and support. We also thank Naama Aviram, Maya Schuldiner and Hugh Pelham for thoughtful discussions, and Sean Munro for critical reading of the manuscript and helpful comments. This work was supported by funding from the US National Institutes of Health (R01 GM085089 to EAM) and the UK Medical Research Council (MC_UP_1201/10 to EAM and MC_UP_1201/8 to WK).

## Author contributions

Conceptualization: EAM. Investigation: NGN, AMC, WK and EAM. Formal analysis: NGN, AMC, JB and EAM. Methodology and Software: JB. Funding acquisition: EAM and WK. Writing – initial draft: NGN, AMC and EAM. Writing – review and editing: NGN, AMC, JB, WK and EAM. Supervision: EAM.

## Abbreviations

COPII: Coat protein II
ERES: 
ER: exit site
sfGFP: Superfolder green fluorescent protein
CLEM: Correlative light and electron microscopy
*bst*: bypass of Sec13
GPI-AP: Glycosylphosphatidylinositol-anchored protein
UPR: Unfolded protein response
FLIP: Fluorescence loss in photobleaching
DTT: dithiothreitol

---

**Figure S1.**
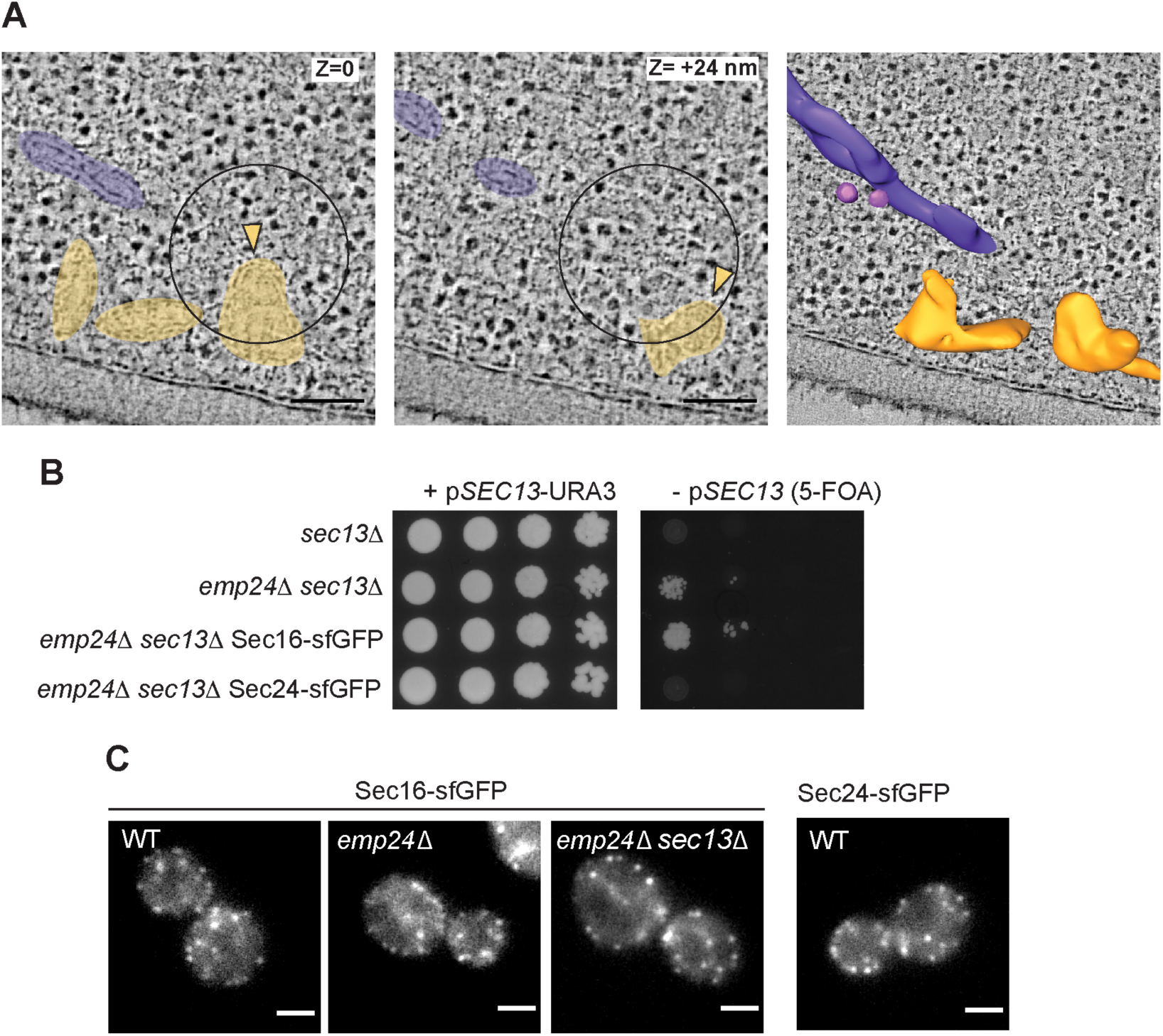
(A) Two virtual tomography slices of a single Sec24-sfGFP positive ERES. Different heights within the correlation are marked (z), showing a multibudded ERES with 2 buds (yellow) and a Golgi complex cisterna (purple). Scale bars are 100 nm. (B) Serial dilutions of the indicated strains were spotted as serial dilutions onto media containing 5-FOA to counter select for the *SEC13-URA3* plasmid and test for viability. On standard media (left panel), all strains grew, whereas growth in the absence of *SEC13* (5-FOA; right panels) was only observed in an *emp24*Δ background. Chromosomal tagging of *SEC16* was tolerated in this background, whereas tagged *SEC24* was not viable. (C) Fluorescence microscopy of the indicated strains expressing *SEC16-sfGFP* and a *SEC24-sfGFP* wild-type strain. Scale bars are 2 µm.

**Figure S2.**
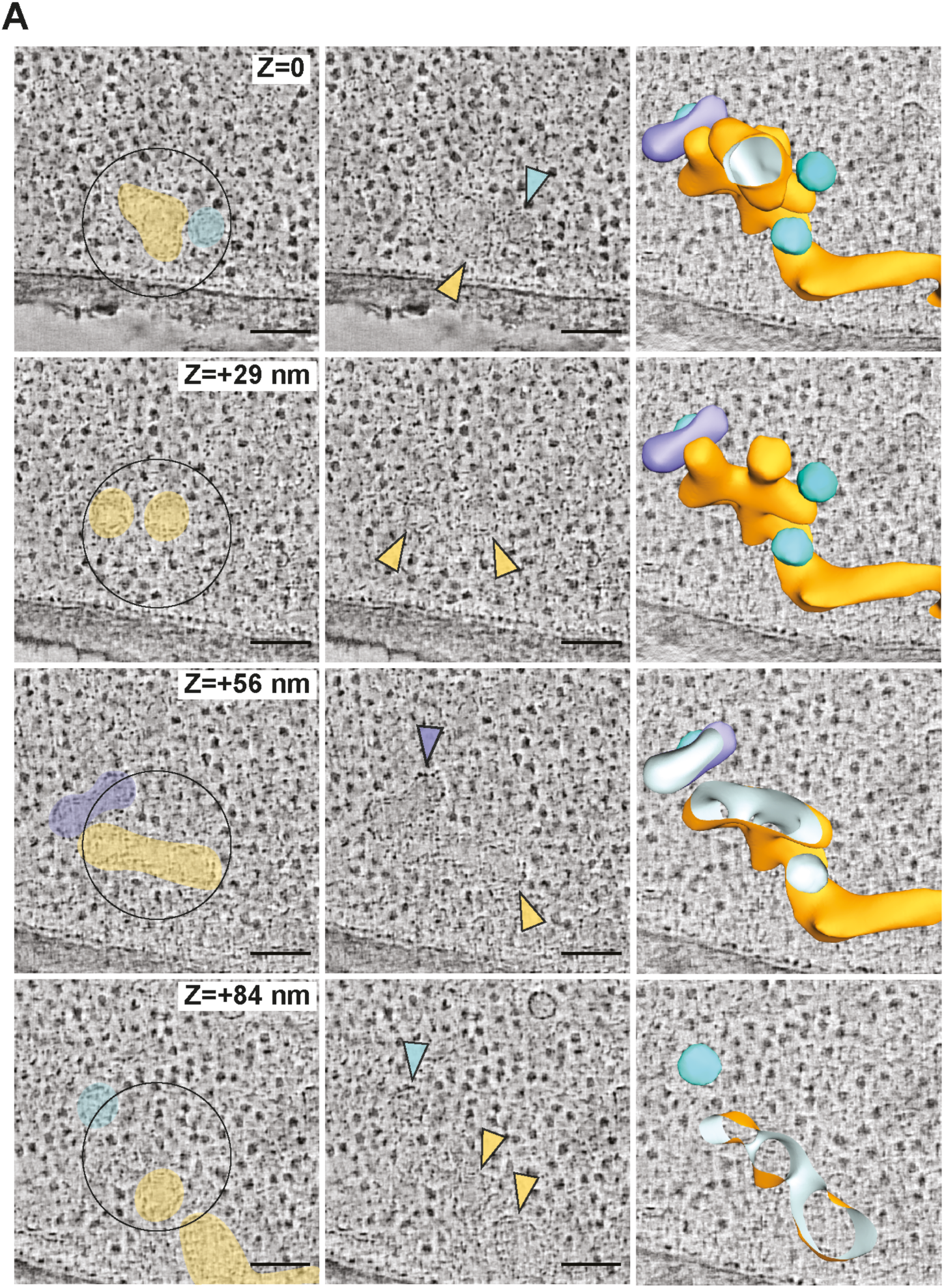
(A) Virtual tomography slices through the 3D volume (z) of an *emp24*Δ *sec13*Δ expressing *SEC16-sfGFP*. Left panels show coloured ER (yellow), vesicles (cyan) and an unidentified tubular compartment (purple). Central panels show the same structures highlighted with coloured arrowheads. Left panels are cut-throughs of a segmentation model of the 3D ultrastructure of membranes at the ERES. Scale bars are 100 nm.

**Figure S3.**
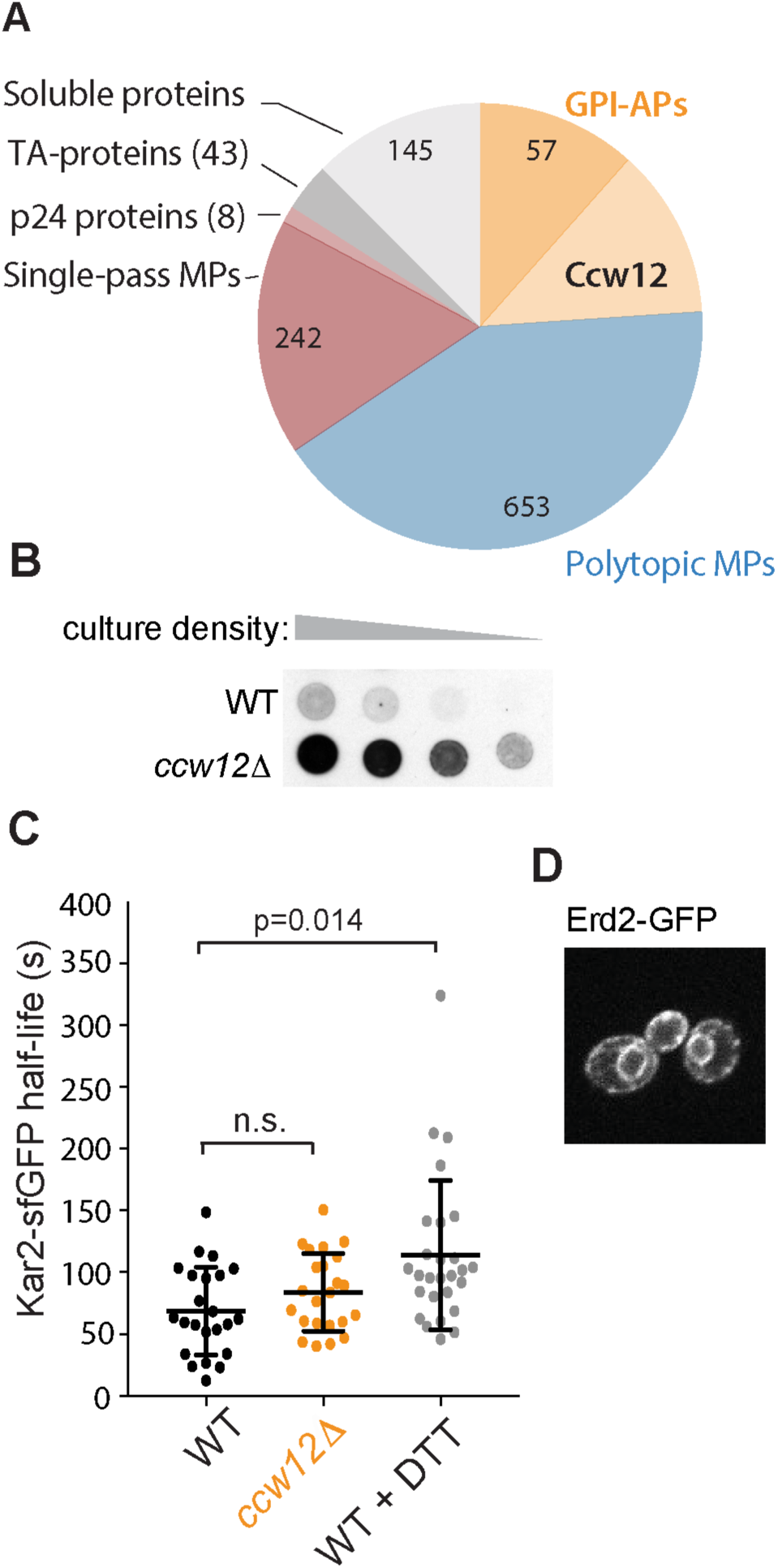
(A) Pie chart of GFP fluorescence of N-terminally tagged secretome proteins (Yofe et al., 2016). Cell wall proteins represent ∼25% of the GFP-secretome, with Ccw12 alone contributing ∼12% of the fluorescent signal. (B) Serial dilutions of wild-type and *ccw12Δ* yeast strains were overlaid with nitrocellulose, and secreted Kar2 detected with Kar2-specific antibodies. (C) Mobility of Kar2-sfGFP was measured by FLIP. Half-time values, calculated as described in figure 3D, of single cells are plotted for the indicated strains or wild-type treated cells with 5 mM DTT for 1 h. Error bars represent SD; n=23. (D) Fluorescence microscopy of *ccw12Δ* cells expressing Erd2-GFP revealed ER localization.

**Figure S4.**
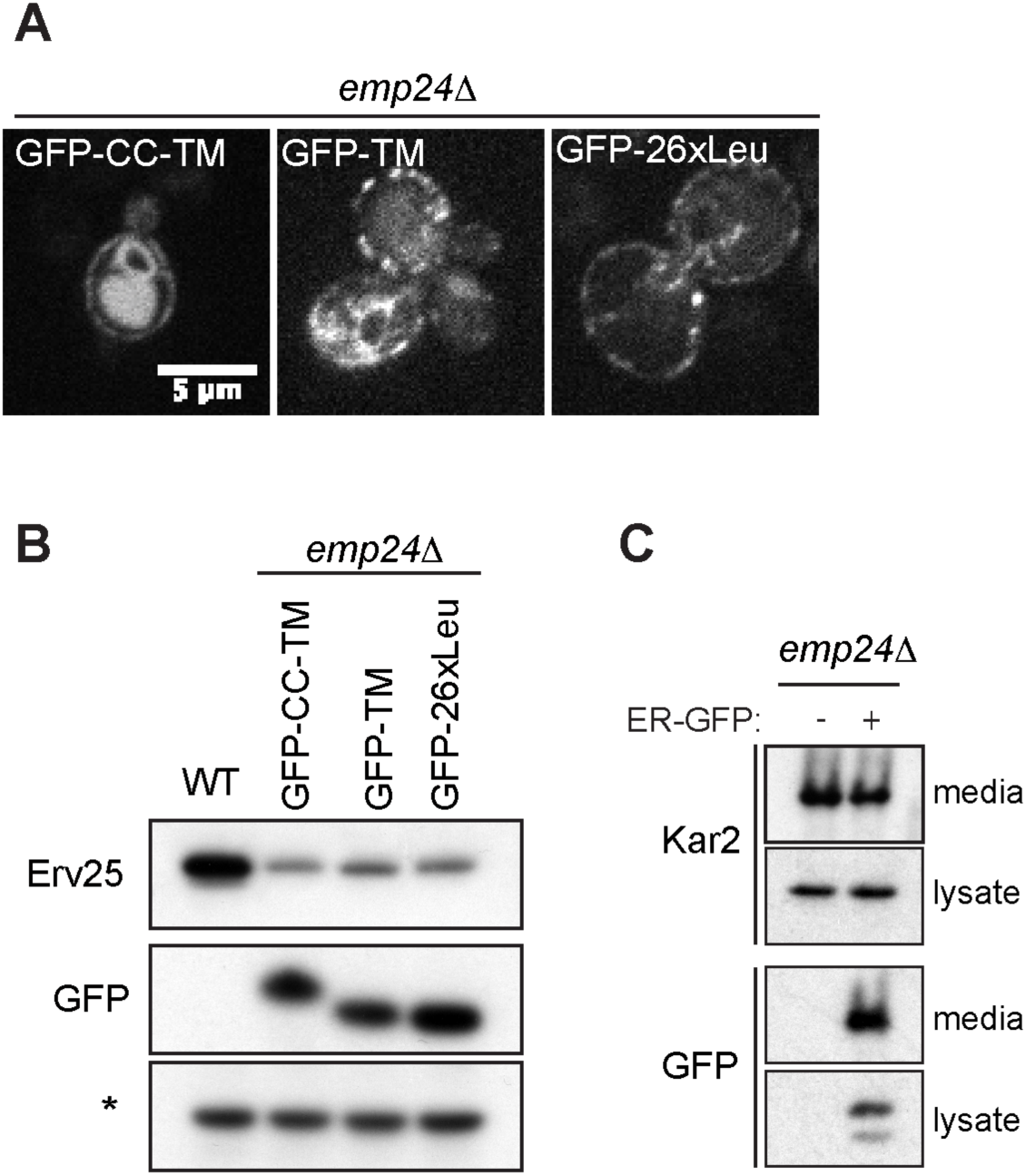
(A) Fluorescence microscopy of *emp24*Δ cells expressing the indicated Emp24 chimeras. (B) Steady state levels of Erv25 in wild-type cells and *emp24*Δ cells expressing the different chimeras indicated were measured from whole cell lysates by immunoblotting using an Erv25-specific antibody. Expression of chimeric proteins was detected from whole cell lysates using an antibody against GFP. A non-specific band labelled with an asterisk is shown as loading control. (C) The amounts of secreted Kar2 and ER-GFP were analysed in *emp24*Δ mutants with and without the ER-GFP plasmid. Secreted proteins in the extracellular media were concentrated using TCA, intracellular proteins were extracted with SDS. Intracellular (lysate) and extracellular (media) proteins were resolved by SDS-PAGE and detected by Western blot against Kar2 and GFP.

