## Supplemental Tables for "Cargo crowding drives sorting stringency in COPII vesicles"

**Table S1: Yeast strains**

| **Strain** | **Description** | **Genotype** | **Source** |
| --- | --- | --- | --- |
| LMY0161 | BY4741 | MATa his3Δ1 leu2Δ0 met15Δ0 ura3Δ0 | Winston et al., 1995 |
| LMY1276 | *Sec24-sfGFP* | MATa his3Δ1 leu2Δ0 met15Δ0 ura3Δ0 SEC24-sfGFP-HIS | This study |
| LMY1277 | *Sec16-sfGFP* | MATa his3Δ1 leu2Δ0 met15Δ0 ura3Δ0 SEC16-sfGFP-KanMX4 | This study |
| LMY1278 | *sec13Δ emp24Δ Sec16-sfGFP* | MATa his3Δ1 leu2Δ0 met15Δ0 ura3Δ0 sec13::HIS emp24::NAT SEC16-sfGFP-KanMX4 | This study |
| LMY1195 | *Sec13D emp24Δ Sec24-sfGFP* | MATa his3Δ1 leu2Δ0 met15Δ0 ura3Δ0 sec13::HIS emp24::NAT SEC24-sfGFP-KanMX4 | This study |
| LMY1279 | *emp24Δ lst1Δ Sec16-sfGFP* | MATa his3Δ1 leu2Δ0 met15Δ0 ura3Δ0 emp24::KanMX4 lst1::NatMX4 SEC16-sfGFP-HIS | This study |
| LMY0438 | *hac1Δ* | MATa his3Δ1 leu2Δ0 met15Δ0 ura3Δ0 hac1::KanMX4 | Euroscarf |
| LMY0426 | *erp1Δ* | MATa his3Δ1 leu2Δ0 met15Δ0 ura3Δ0 erp1::KanMX4 | Euroscarf |
| LMY0375 | *erp2Δ* | MATa his3Δ1 leu2Δ0 met15Δ0 ura3Δ0 erp2::KanMX4 | Euroscarf |
| LMY0573 | *erp1Δ hac1Δ* | MAT? his3Δ1 leu2Δ0 met15? ura3Δ0 lys2? erp1::KanMX4 hac1::NatMX | Copic et al., 2009 |
| LMY0574 | *erp2Δ hac1Δ* | MAT? his3Δ1 leu2Δ0 met15? ura3Δ0 lys2? erp2::KanMX4 hac1::NatMX | Copic et al., 2009 |
| LMY0394 | *emp24Δ* | MATa his3Δ1 leu2Δ0 met15Δ0 ura3Δ0 emp24::KanMX4 | Euroscarf |
| LMY1250 | *ccw12Δ* | MATa his3Δ1 leu2Δ0 met15Δ0 ura3Δ0 ccw12::KanMX4 | Euroscarf |
| LMY1251 | *UPRE-KAR2* | MATa ura3-1 leu2-3 his3-11 trp1-1 can1-100 ade2-1 ura3-52::UPRE-KAR2-URA3 kar2::KanM | Chia-Ling et al., 2012 |
| LMY1252 | *UPRE-KAR2 emp24Δ* | MATa ura3-1 leu2-3 his3-11 trp1-1 can1-100 ade2-1 ura3-52::UPRE-KAR2-URA3 kar2::KanMX emp24::LEU2 | This study |
| LMY1253 | *upred-KAR2* | MATa ura3-1 leu2-3 his3-11 trp1-1 can1-100 ade2-1 ura3-52::upred-KAR2-URA3 kar2::KanMX | Chia-Ling et al., 2012 |
| LMY1254 | *upred-KAR2 emp24Δ* | MATa ura3-1 leu2-3 his3-11 trp1-1 can1-100 ade2-1 ura3-52::upred-KAR2-URA3 kar2::KanMXemp24::LEU2 | This study |
| LMY1255 | *kar2-1* | MATa ura3-52 leu2-3 ade2-101 kar2-1 | Kabani et al. 2003 |
| LMY1256 | *kar2-1 emp24Δ* | MATa ura3-52 leu2-3 ade2-101 kar2-1 emp24::KanMX4 | This study |
| LMY1081 | *lst1Δ* | MATa his3Δ1 leu2Δ0 met15Δ0 ura3Δ0 lst1::NatMX4 | D’Arcangelo et al., 2015 |
| LMY1257 | *emp24Δ lst1Δ* | MATa his3Δ1 leu2Δ0 met15Δ0 ura3Δ0 lst1::NatMX4 emp24::KanMX4 | This study |
| LMY1258 | *ccw12Δ lst1Δ* | MATa his3Δ1 leu2Δ0 met15Δ0 ura3Δ0 ccw12::KanMX4 lst1::LEU2 | This study |
| LMY1271 | *Kar2-sfGFP* | MATa his3Δ1 leu2Δ0 met15Δ0 ura3Δ0 Kar2-sfGFP-HDEL-*NatMX4* | This study |
| LMY1272 | *emp24Δ Kar2-sfGFP* | MATa his3Δ1 leu2Δ0 met15Δ0 ura3Δ0 emp24::KanMX4 Kar2-sfGFP-HDEL-*NatMX4* | This study |
| LMY1273 | *ccw12Δ Kar2-sfGFP* | MATa his3Δ1 leu2Δ0 met15Δ0 ura3Δ0 ccw12::KanMX4 Kar2-sfGFP-HDEL-*NatMX4* | This study |
| LMY0804 | *emp24Δ Erd2-GFP* | MATa his3Δ200 leu2Δ1 ura3-52 lys2-801 am emp24::KanMX4 Erd2-GFP-*LEU2* | Copic et al., 2009 |
| LMY1275 | *ccw12Δ Erd2-GFP* | MAT? his3Δ1 leu2Δ0 met15? ura3Δ0 lys2? ccw12::KanMX4 Erd2-GFP-*LEU2* | This study |

**Table S2: Plasmids**

| **Plasmid** | **Description** | **Source** |
| --- | --- | --- |
| *pGAL-CPY*-HA* | pTS210-GAL1-CPY*-HA | Kawaguchi et al., 2010 |
| *pGAL-CPY*-Δ1-HA* | pTS210-GAL1-CPY*-D1-HA | Kawaguchi et al., 2010 |
| *pSP-GFP-CCTM* | pRS316-*GAL1-SP-GFP-CoiledCoil-TMD- Emp24 cytosolic domain* | GeneScript (This study) |
| *pSP-GFP-TM* | pRS316-*GAL1-SP-GFP-TMD- Emp24 cytosolic domain* | GeneScript (This study) |
| *pSP-GFP-26xLeu* | pRS316-*GAL1-SP-GFP-26xLeu-Emp24 cytosolic domain* | This study |
| *pSP-FLAG-Cp* | p426-*GAL1-SP-FLAG-Capsid protease (Cp) domain* | This study |
| *pER-GFP* | pRS415-ss-sfGFP-Myc | Snapp et al., 2006 |
| *pEmp24-sfGFP* | YCplac111-*EMP24-sfGFP* | D'Arcangelo et al., 2015 |
| *pLST1* | pRS313-*LST1* | D'Arcangelo et al., 2015 |
| *pLST1-B* | pRS313-*lst1*−b (K543M,R545M) | D'Arcangelo et al., 2015 |
